# A protein microarray-based *in vitro* transglutaminase assay platform for epitope mapping and immunogen design: implication for transglutaminase-mediated immunodominant determination

**DOI:** 10.1101/2021.05.12.443622

**Authors:** Chen Liu

## Abstract

Transglutaminases (TGs) are a family of crosslinking enzymes catalyzing the formation of intra- and inter-molecular glutamine-lysine isopeptide bonds in a calcium dependent manner. Activation of transglutaminases is pathogenically associated with severe human diseases including neurodegenerations, cardiovascular diseases, and autoimmune diseases. Although continuous efforts determining the enzymes’ substrate preference have been witnessed, a high-throughput assay platform with the “omic” efficiency is still missing for the global identification of substrate-specific TG modification sites. In this study we report a protein microarray-based *in vitro* TG assay platform for rapid and large-scale (up to 30000 reactions per chip) determination of the glutamine (Q)-bearing TG modification motifs. With this platform we identified the Q16 in superoxide dismutase 1 and Q109 in alpha-synuclein as the primary modification sites for tissue transglutaminase (TG2), the most ubiquitous member of the enzyme family. Of particular interest, we found a close match between the modification motifs and published vaccine epitope sequences in alpha-synuclein, implying an essential and intrinsic role transglutaminase might play in the determination of immunodominant epitopes. Our data collectively suggest the glutamine and its follow-up five residues on the C terminal of a protein compose a minimal determinant motif for TG2 modification and, more importantly, the TG2 modification motifs determined by our platform could finally correspond to the substrate’s immunodominant epitope sequences in antigen processing. To establish an efficient approach to optimize the enzyme modification motifs and screen for site-specific interfering peptides, we scanned the known TG2 modification motifs with *onchip* amino-acid swapping and glutamine repeat addition, and obtained multiple variants with significantly upregulated TG2 reactivity. This approach could also be employed to improve the target peptide’s immunogenicity. Taken together, our synthetic transglutaminase assay platform might be able to deliver a precise epitope blueprint for immunotherapeutic targeting and provide pilot and directional studies for TG-based peptide discovery and immunogen design.

## Introduction

Transglutaminases (TGs) in mammalian cells are a family of crosslinking enzymes posttranslationally catalyzing the covalent isopeptide bond formation between the γ-carboxamide groups of a peptide-bound glutamine residue and a free amine or a peptide-bound lysine in a calcium dependent manner^1–4^. Functionally, TG-mediated isopeptide modification results in the stabilization and aggregation of substrate proteins by facilitating the assembly of supramolecular structure resistant to proteolysis^1, 5^. Activation of transglutaminase and TG-mediated isopeptide modification have been demonstrated to be pathogenically associated with severe human diseases including neurodegenerations^6–9^, cardiovascular diseases^10–13^, and autoimmune diseases^12, 14, 15^. Accumulation of pathogenic proteins crosslinked by transglutaminase has long been recognized as a hallmark of major neurodegenerative disorders including Alzheimer’s disease (AD)^16–21^, Parkinson’s disease (PD)^22, 23^, and Huntington’s disease (HD)^24, 25^. In cardiovascular diseases, transglutaminases contribute to the formation of atherosclerotic plaques by crosslinking extracellular matrix (ECM) proteins^26–31^ and to hypertensive disorders by modifying and sensitizing vasopressor receptors^32, 33^. Transglutaminases are also well-known to participate in autoimmune responses through posttranslational modifications favoring the generation of neoantigens^34–37^. Successful amelioration of key disease features by transglutaminase inhibitors in the pre-clinical animal models of related disorders not only establishes TG as an effective therapeutic target but also potentiates transglutaminase inhibitors as useful drugs for the disease treatment^38–42^. However, diagnosis of the related complications in patients usually accompanies a significant build-up of TG-crosslinked aggregates or plaques, and simply inhibiting the enzymes’ activity is not sufficient to clean the build-ups. Given this concern, characterizing substrate-specific crosslinking sites and understanding the sequence preference of transglutaminase substrate would be of great importance in designing immunotherapeutic strategies capable of cleaning the aggregates to revoke the disease progression.

To determine the enzymes’ substrate preference and map the modification sites among various protein substrates, continued efforts using phage display^43–45^, mass spectrometry^46–49^, protein arrays^50, 51^, and bioinformatics tools^52–54^ have been witnessed. Although these studies greatly advanced our knowledge regarding the modification patterns of transglutaminases, a high-throughput platform with the efficiency of systems biology is still missing for the identification of substrate-specific TG modification sites. In this study, by combining the tagged amine donor dansyl-cadaverine-based *in vitro* TG assay^55^ with a protein microarray^56^ we report a platform for rapid and large-scale (up to 30000 reactions per chip) determination of the glutamine (Q)-bearing TG modification motifs. We tested the platform with peptides from neurodegenerative proteins including alpha-synuclein and superoxide dismutase 1 and mapped their primary modification sites for tissue transglutaminase (TG2), the most ubiquitous member of the enzyme family^1–4^. To our surprise, the modification motifs in alpha-synuclein match well the published immunodominant epitope sequences obtained from the full-length protein immunization. Our data further indicate the glutamine and its follow-up five residues on its C terminal compose a minimal determinant motif for TG2 modification that could finally become core part of the substrate’s epitope sequences in antigen processing. To manipulate the TG2 modifications on a certain protein and screen for site-specific interfering peptides, we employed *onchip* amino-acid scanning ^57,58^ and glutamine repeat addition methods for the optimization of modification motifs. By scanning the TG2 modification motif QQIV in the extracellular matrix protein fibronectin, we confirmed the platform’s capability to serve TG-based peptide discovery and immunogen engineering.

## Methods

### *Onchip in vitro* TG2 assay

To identify the glutamine residues that can be modified by tissue transglutaminase on the peptide microchip, the synthesized peptide microchip was incubated with 100 ug/ml guinea pig liver tissue transglutaminase (Sigma) and 3 mM dansyl-cadaverine (Sigma) in 1 ml of TBS containing 5 mM Calcium Chloride and 1 mM DTT at 37 degree for 30 minutes. Afterwards, the peptide chip was washed at least 3 times with TBS. After washing off tissue transglutaminase and dansyl-cadaverine molecules bound on the synthesized peptides, the dansyl-cadaverine conjugated on the chip was tracked by rabbit anti-dansyl antibody (Invitrogen) followed by Alexa Fluor 594-labeled anti-rabbit secondary antibody (Invitrogen). Fluorescent microchip figures were quantified and analyzed with ArrayPro32. Original array figures and data are available upon request.

## Results

### Mapping TG2 modification sites in neurodegenerative proteins with high-throughput *in vitro* assay platform

To establish a high-throughput assay platform for the rapid and large-scale identification of TG2 modification sites in disease-related proteins, we synthesized on microchips the glutamine-bearing motifs in superoxide dismutase 1 (SOD1) and alpha-synuclein, the pathogenic proteins in amyotrophic lateral sclerosis (ALS) and Parkinson’s disease, respectively.

To characterize the glutamine-bearing TG2 modification motifs on these synthesized peptides, dansyl-cadaverine, a well-established amine donor in transglutaminase reaction, was covalently conjugated to the glutamine residues on the peptide chip by purified TG2 with the help of calcium (Figure 1A). After washing off tissue transglutaminase and dansyl-cadaverine molecules bound on the synthesized peptides, the dansyl-cadaverine conjugated on the chip was probed by anti-dansyl antibody followed by Alexa Fluor 594-labeled anti-rabbit secondary antibody. In this way the level of dansyl-cadaverine incorporation on a certain peptide was measured by the fluorescent intensity.

**Figure 1.**
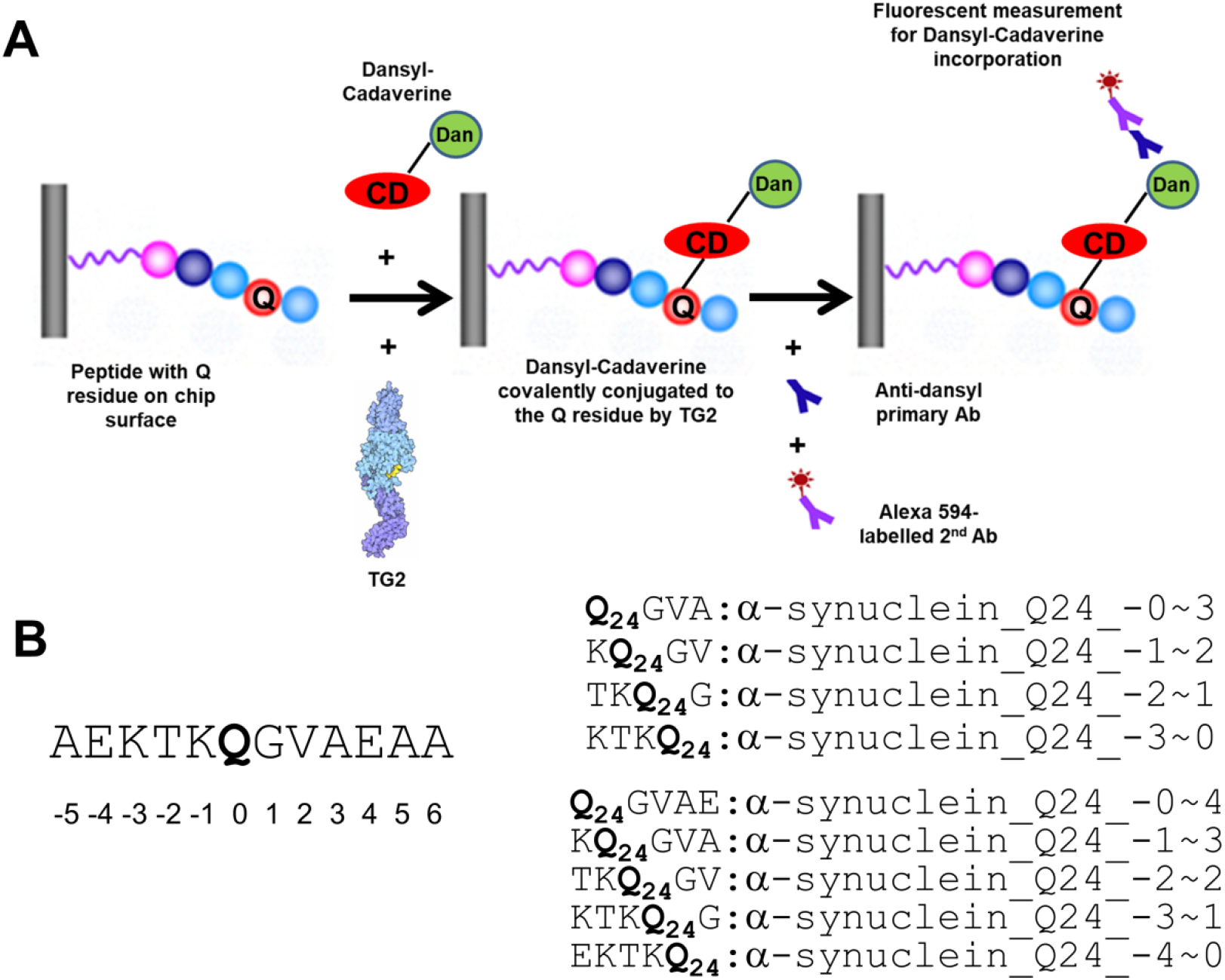
Peptide microarray-based *in vitro* TG2 assay platform for rapid and high-throughput identification of modification Q sites among proteins of interest. **A.** Assay flow chart for the Q mapping platform (Q: glutamine; CD: cadaverine; Dan: dansyl; Ab: antibody; TG2: tissue transglutaminase). **B.** Peptide layout strategy and naming rule on the microchip using the first glutamine (Q24) motif in human alpha-synuclein as the example. Each peptide where position of Q is defined as 0 and the one directly before or after it as −1 or 1 is named as *protein name_Q residue# in the entire protein*_*N terminal residue# in the peptide~C terminal residue# in the peptide*.

Each peptide synthesized on the chip has a length of at least 4 amino acids, and its maximum length could be up to 12 mer. And the glutamine residue needs to appear in each position of the peptide once to ensure the thorough coverage of the screening (Figure 1B). In this way the peptide screening may also pattern the substrate sequence optimal for the modification. For example, in human alpha-synuclein the surrounding sequence of the first glutamine is AEKTKQ24GVAEAA (Figure 1B). So the sequences for its 4 mer peptides would be QGVA, KQGV, TKQG, and KTKQ, and those for 5 mer would be QGVAE, KQGVA, TKQGV, KTKQG, and EKTKQ. Up to 11 residues on either N or C side of the glutamine is covered on the chip. Each peptide is named as *protein name_Q residue# in the protein_N terminal residue# in the peptide~C terminal residue# in the peptide (In the peptide position of Q is defined as 0, and the one directly before or after it as −1 or 1).* So the 4 mer peptide KQ_24_GV is named as α-synuclein_Q24_-1~2 (Figure 1B). Therefore, for each glutamine residue in a given protein, the initial number of peptide variants synthesized on the chip will be 4+5+6+7+8+9+10+11+12=72. The corresponding peptides with Q to S swap are also synthesized on the same chip as negative controls.

With this approach we identified Q16 in superoxide dismutase 1 and Q109 in alpha-synuclein as the primary modification sites for TG2 (Figure 2B and 2C). In human SOD1 protein there are 3 glutamine residues including Q16, Q23, and Q154. Compared with Q23 and Q154 counterparts, the Q16 peptides with the 5 mer motif directly following the glutamine residue (QGIINF) showed significantly higher fluorescent intensities (>6000), and the Q to S swap could effectively reduce their fluorescent levels, indicating the Q16 residue is the most probable TG2 modification site in the protein (Figure 2B). Similarly, among the 6 glutamine residues in human alpha-synuclein protein, the Q109 residue with its follow-up 5 mer motif (QEGILE) elicited the strongest dansyl-cadaverine incorporation signal and thereby was identified as the TG2 modification site of the protein (Figure 2C). Aligned data with top hits from SOD1 and alpha-synuclein collectively suggest that the Q and its follow-up 5 residues compose a minimal determinant motif for TG2 modification (Figure 2D), which is further confirmed by the peptides with truncated minimal determinant motifs (not shown).

**Figure 2.**
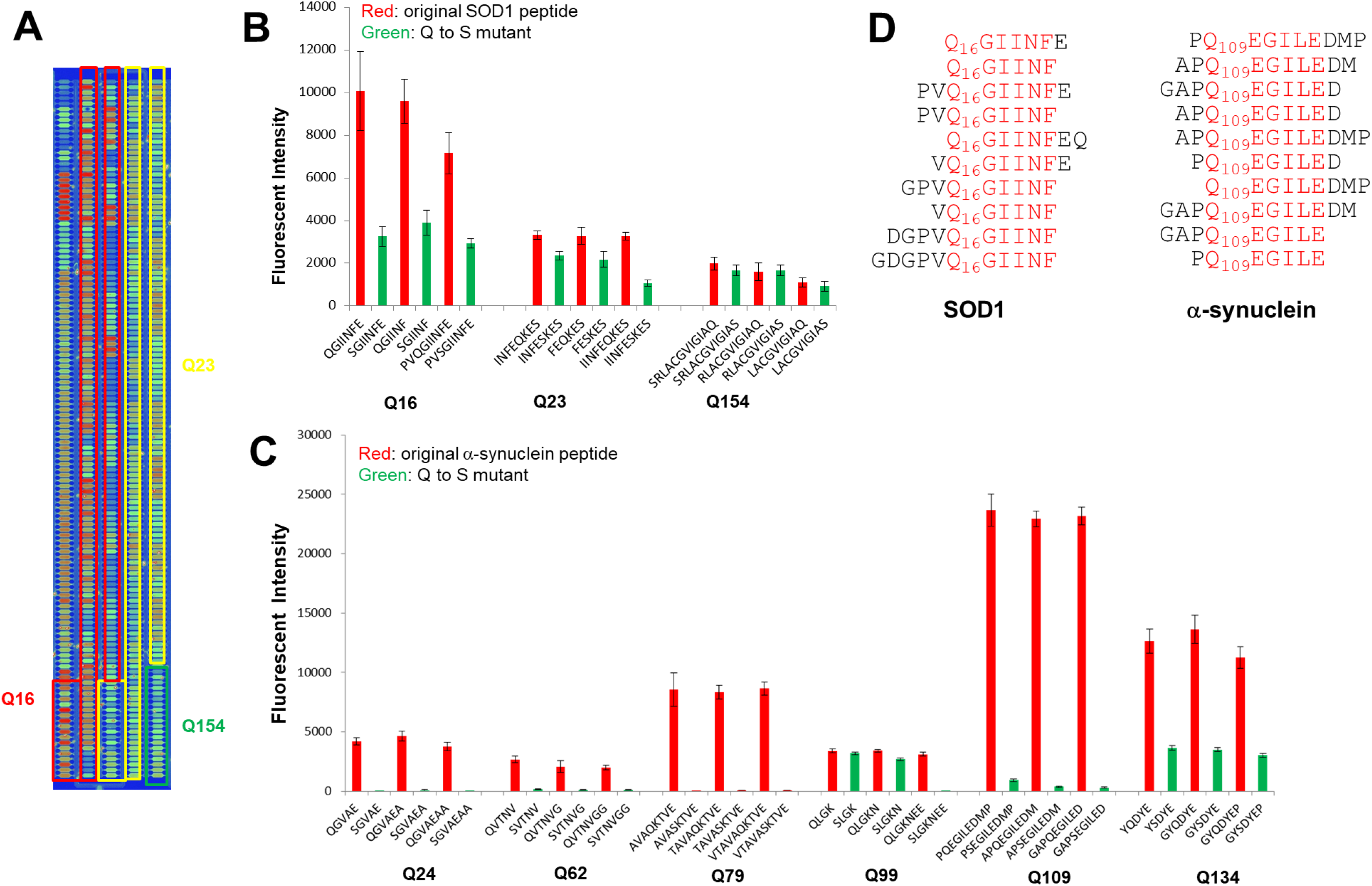
Identification of the glutamine-bearing TG2 modification sites in SOD1 and alpha-synuclein with the high-throughput *in vitro* assay platform. **A.** Representative image of the protein microarray-based *in vitro* TG2 assay with the Q peptides in SOD1 showing that the first Q (Q16) peptides (red box) have a generally stronger reaction signal (red box) than the other two (Q23:yellow box; Q154:green box). Q16 residue in SOD1 (**B**) and Q109 in alpha-synuclein (**C**) are identified as the primary TG2 modification sites (p<0.05 vs other residues or mutant controls in **B** and **C**; n=triplicates; data=mean ± SEM). The top 3 peptide hits of each Q residue in the TG2 reaction are plotted together with their Q to S mutants. **D.** Alignment of the top 10 hits of Q16 peptides in SOD1 and Q109 peptides in alpha-synuclein indicate their minimal determinant motifs.

### Optimizing TG2 modification sites with the *onchip* amino-acid scanning

The small peptide QQIV is a transglutaminase substrate identified in the extracellular matrix protein fibronectin. As an amine acceptor, the peptide has been demonstrated to be an effective competitive inhibitor for the transglutaminase reaction^59^. Given this, we chose it as one of the positive substrate peptides on the peptide chip. On our assayed chip the QQIV peptide showed a fluorescent intensity of ~5000 which is much higher than those (~800) of four-residue negative control peptides without glutamine residues and the chip background (Figure 3A). These results confirmed the efficiency and specificity of the dansyl-cadaverine incorporation in our assay system. To scan for a more preferred TG2 substrate and thereby for a more optimized competitive inhibitor than QQIV, we swapped the isoleucine (I) residue in this small peptide to every other amino acid. Except one variant with reduced transglutaminase reactivity, 14 out of the 19 swapped peptides show significantly higher fluorescent signals and the improvement could be up to 3 folds (Figure 3A), suggesting their better candidacy for competitive inhibitor and substrate of TG2 modification than the original QQIV. However, transglutaminase reactivity of most of these 14 mutants was significantly compromised when the hydrophobic V residue was changed to the hydrophilic G (data not shown). To further test the amino acid-scanning approach with the minimal determinant motif for TG2 modification, we swapped to any other amino acid the first leucine residue in the 12-mer small peptide sequence REQLYLDYNVFS, a known TG2 substrate found in a phage display library. Through pan-amino acid scanning at the L residue we found 6 variants (L to N, S, E, R, V or T) with significantly higher transglutaminase reactivity and 6 (L to G, K, M, W, Y or F) with lower reactivity (Figure 3B). Taken together, our result indicates that the residues within the minimal determinant motif could be reasonable targets for the optimization of modification site and the design of substrate-specific interfering peptides.

**Figure 3.**
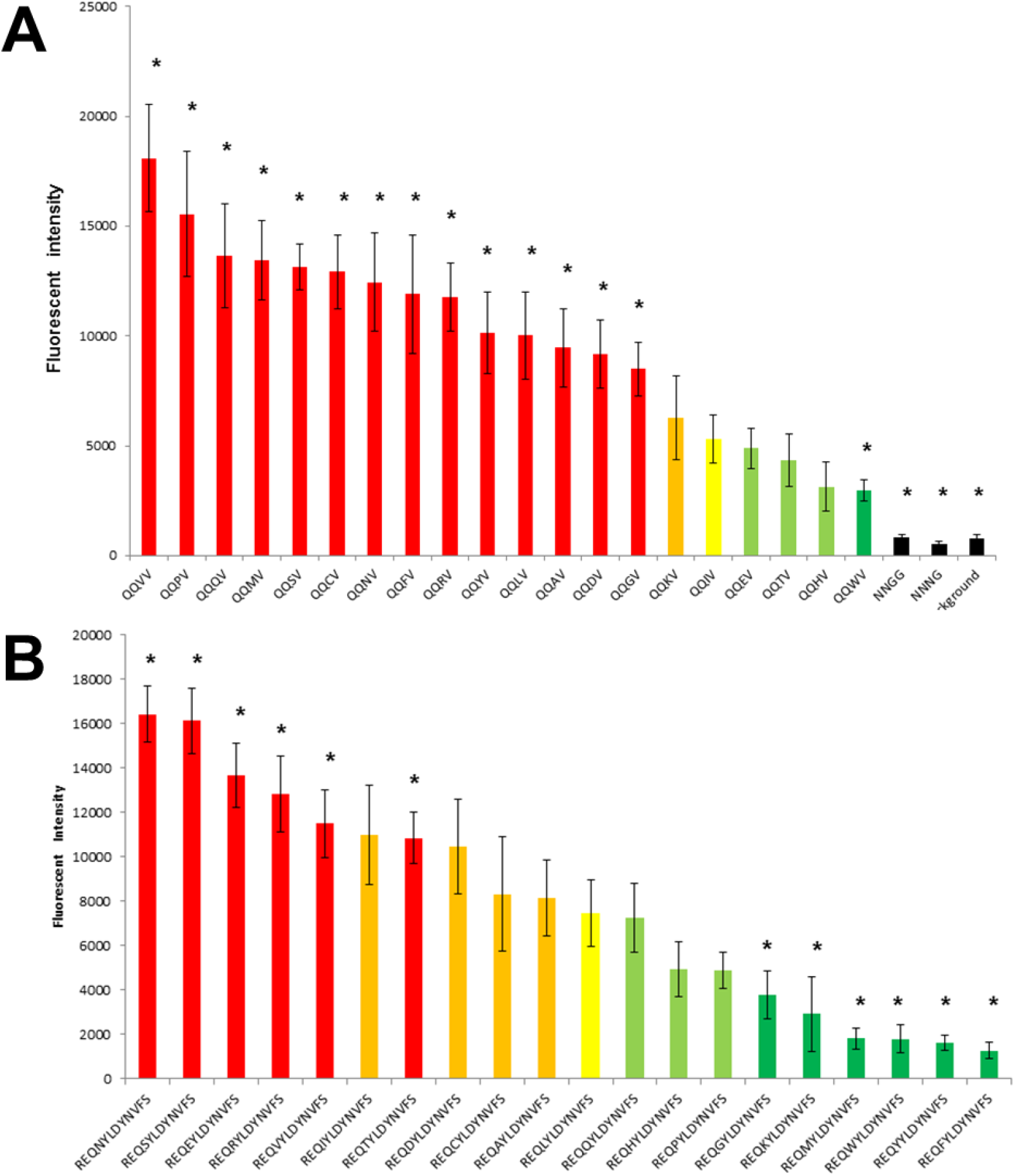
*Onchip* amino-acid scanning generates peptides with significantly changed reactivity with TG2. **A.** Pan-amino acid swapping at the I site of the peptide sequence QQIV generates 14 variants (red bars) with significantly higher TG2 reactivity and 1 lower (dark green) (*P<0.05 versus QQIV; n=triplicates; data=mean ± SEM). **B.** Amino acid scanning at the L residue directly after the Q site of the TG2 substrate peptide REQLYLDYNVFS also obtained variants with significantly changed TG2 reactivity (*P<0.05 versus REQLYLDYNVFS; n=triplicates; data=mean ± SEM).

### Elevating peptide’s reactivity with transglutaminase by adding glutamine repeats

Characterized in neurodegenerative complications like the polyglutamine diseases, glutamine repeats elicit excellent TG substrate properties in polypeptides by functioning as efficient amine acceptors, and thereby are considered as a biochemical cause for the pathogenesis. To test this feature on our platform, we randomly added the double glutamine repeat motif QQXX (X stands for any of the 20 amino acids) at the N terminal of the peptide QQIV. Consistent with previous findings, the addition of double glutamine repeat motif resulted in a significant upregulation of the TG2-mediated conjugation of dansyl-cadaverine as the average fluorescent intensity (~10000) of the 400 QQXXQQIV peptides is two folds higher than that (~5000) of the original QQIV peptide (Figure 4A) and 297 out of the 400 QQXXQQIV peptides show a fluorescent intensity of more than 5000 on our assayed chip (Figure 4B). With an average fluorescent intensity of ~20000 (Figure 4A), the top 40 QQXXQQIV peptides could serve as competitive inhibitors to block TG2’s modification on the QQIV motif of fibronectin. And among them, the peptides with P at the third residue or I at the fourth show up at most times (Figure 4C), indicating a preferred pattern for the linker region between glutamine repeats.

**Figure 4.**
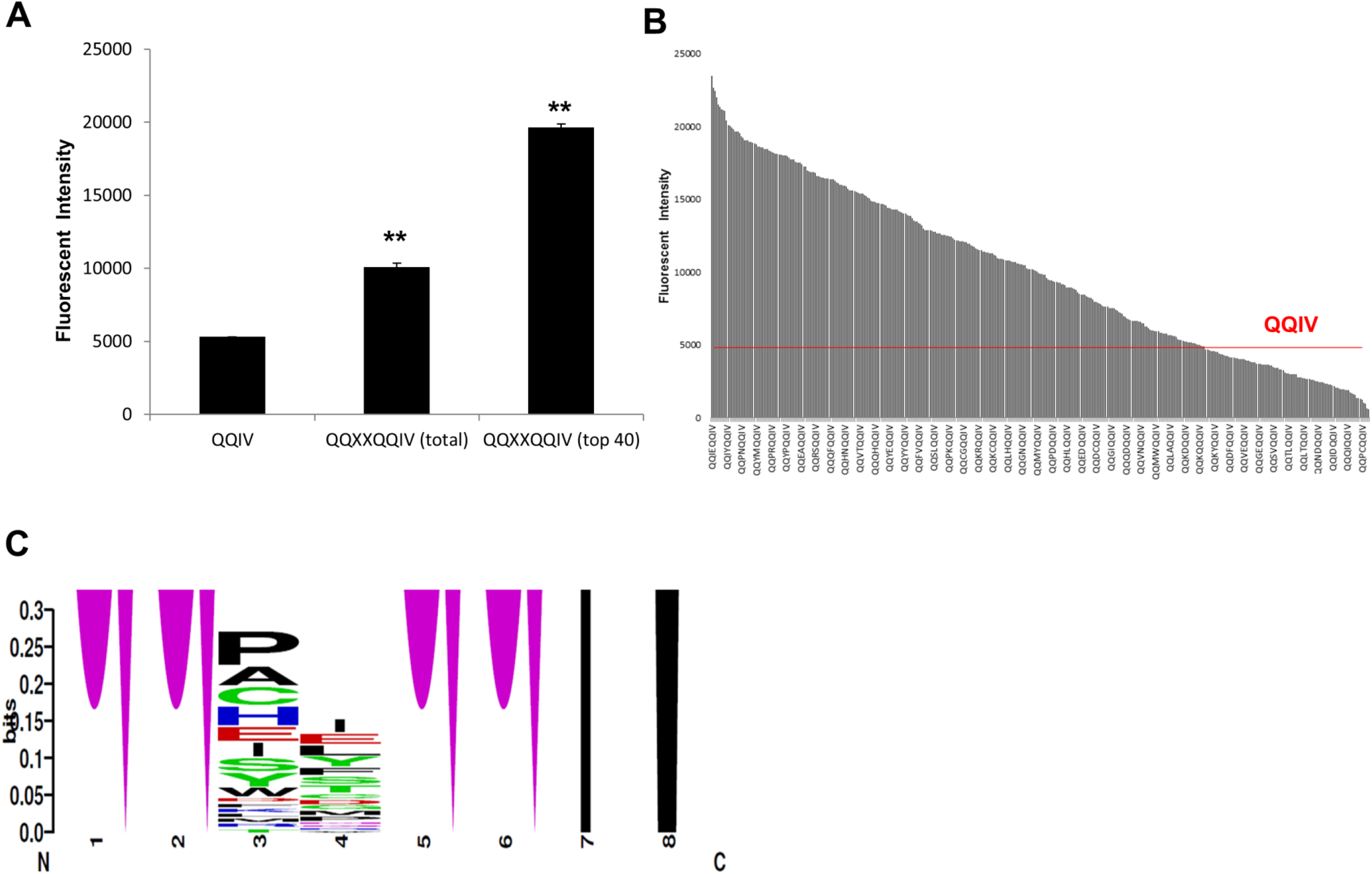
Addition of glutamine repeat elevates TG2 reactivity of the fibronectin peptide QQIV. **A.** Addition of glutamine repeat at the N terminal of the fibronectin peptide QQIV results in significant increase in dansyl-cadaverine incorporation as measured by fluorescent intensity on the chip (**P<0.01 versus QQIV; X=any of 20 amino acids; n=triplicates; data=mean ± SEM). **B.** Majority (297 out of total 400) of the QQXXQQIV variants show a higher level of TG2 reactivity as measured by fluorescent intensity on the chip (the fluorescent intensity of QQIV is ~5000 as indicated). **C.** Among the top 40 QQXXQQIV peptides with the highest fluorescent intensity, the peptides with a P at the third residue or I fourth appear most.

## Discussion

In this study we report a protein microarray-based *in vitro* TG assay platform for fast and high throughput identification of the glutamine (Q)-bearing TG modification motifs among proteins of interest. With this platform we characterized the Q16 in superoxide dismutase 1 and Q109 in alpha-synuclein as the primary TG2 modification sites. Our data indicate the glutamine and its follow-up five residues make up a minimal determinant motif for TG2 modification. To optimize the site for TG2 modification we used an *onchip* amino-acid scanning method in which the residues near the glutamine residue are swapped with any other amino acid to manipulate its modification level. Using this approach we scanned the isoleucine residue in TG2 modification motif QQIV of fibronectin and obtained 14 variants with significantly higher TG2 reactivity and 1 with lower reactivity. We further confirmed this approach by scanning the first leucine residue within the minimal determinant motif of the 12-mer peptide REQLYLDYNVFS. Consistent with the findings from polyglutamine diseases, we also showed that the addition of glutamine repeats with optimized linker residues are able to significantly elevate the peptide QQIV’s transglutaminase reactivity.

Protein aggregation is a hallmark of neurodegenerative diseases including but not limited to Alzheimer’s disease, Huntington’s disease, and Parkinson’s disease, where TG2-mediated crosslinking of the aggregates has been consistently characterized. Moreover, by preventing substrates’ ubiquitination^60^, TG2 crosslinking could directly impair the targeted degradation of the aggregates and is thus considered to play a pathogenic role. To map the crosslinking sites for the development of immunotherapies targeting the pathogenic aggregates, we designed a protein microarray-based *in vitro* TG2 assay platform and confirmed its efficacy by identifying the TG2 modification sites in alpha-synuclein and superoxide dismutase 1. With this approach the Q109 in alpha-synuclein is characterized as the primary glutamine site for the enzyme modification. Other residues near the c-terminus including Q79 and Q134 are also identified as modification sites. Of particular interest, these modification motifs correspond to the epitope sequences found in animals with full-length protein immunization^61, 62^, in which antibodies recognize aa85-99, aa109-123, aa112-126, and aa126-138; B cells aa106-125; and T cells aa76-95 and aa106-125. The motifs with the primary modification site Q109 are the only type of glutamine motifs present among all the antibody, B and T cell epitopes. Consistent with the roles of transglutaminase in the generation of autoimmune neoantigens^34–36, 63^, these findings suggest a previously overlooked essential step mediated by transglutaminase for the determination of immunodominant epitopes in the endogenous antigen processing and presentation pathways. Given the presence of glutamine residues, an involvement of transglutaminase crosslinking will fit into the established models of immunodominance including hen-egg lysozyme (HEL) aa46-61^64, 65^. Immunodominance refers to the preference of immunological responses for only a small subset of the peptides generated in the processing of a certain antigen protein. Origination of antigen peptides requires ubiquitin labelling of substrate proteins for their transportation to the follow-up proteolytic digestion in either proteasome or lysosome. Thus, the epitope determination within an antigen peptide would first require its proteasomal^66, 67^ co or lysosomal^68^ survival in antigen processing. Transglutaminase crosslinking is known to render its substrates proteolytic resistance^1^. And the glutamine-lysine isopeptide modification introduced by the crosslinking enzyme is believed to prevent ubiquitination of substrate proteins via depleting lysine availability^60^. These two machineries together would ensure the modified peptides to emerge as the candidates of antigenic determinants. More importantly, the glutamine-lysine isopeptide modification over the peptides could serve as a crucial tag of determinants for their recognition by MHC and/or TCR-pMHC complex in antigen presentation. To induce a decent immune response, the determinant peptides need to bind the MHC at the greater than threshold affinity^69^. The side chain with isopeptide bond might provide better anchorage through unconventional chemical interaction with MHC residues at the interface^70^. Interestingly, the length of glutamine motifs with the highest transglutaminase reactivity in alpha-synuclein on the microchip matches the size of class I MHC binding peptides (8-10mer). However, the identification of MHC bound peptides without glutamine and lysine residues^71^ implies that the dynamic interaction and positioning of the isopeptide side chain in the TCR-pMHC complex could be the real rate-limiting step in the immunodominant determination. Based on this rationale, the TG2 modification motif determined by our assay platform should directly correspond to the substrate’s immunodominant epitope sequence recognized by the immunity, and our platform could be a general system applying to most epitope mapping efforts.

Amyotrophic lateral sclerosis is associated with SOD1 mutations that may greatly alter the protein’s reactivity with transglutaminase and thereby contribute to the protein aggregation. With the synthesized mutant peptides on chip, our platform is also able to determine the mutation-specific epitopes through their unique TG2 modification barcodes for immunotherapeutic targeting. To a greater extent, by measuring the mutation-related crosslinking signatures our synthetic assay platform could map the disease-specific epitopes for the disorders frequently associated with genetic mutations like cancer.

In this study, through scanning the residue directly following glutamine of the minimal determinant motif in the peptides QQIV and REQLYLDYNVFS, we obtained variants with significantly changed transglutaminase reactivity. Inspired by polyglutamine diseases, we also introduced into the fibronectin peptide QQIV additional glutamine repeats with optimized adaptor sequence. With either of the approaches, we obtained variants with significantly elevated transglutaminase reactivity that are supposed to be associated with better immunogenicity. Therefore, our assay platform would not only deliver candidates for site-specific competitive inhibitors, but also serve as a dynamic immunodominant epitope design tool overcoming the high mutation rate, genetic polymorphism, and low immunogenicity in therapeutic targets. For instance, mutations help viruses evade immune surveillance generated by the conventional vaccines, and compromised immunogenicity in conserved viral domains exacerbates the case. Following a global determination of immunodominant epitopes, improved and overlapping immunogens covering most mutational possibilities or less immunodominant but more conserved domains could be designed with the *onchip* positional scanning or glutamine introduction approach on the *in vitro* transglutaminase assay platform for next-generation universal vaccines with spatiotemporal coverage. In this way, a universal, straightforward, and convenient approach for immunodominant epitope determination and optimization could be established for next-generation immunotherapeutics treating most human diseases.

## Supporting information

Synuclein TG array figure

Synuclein TG array data

Summarized TG array data

## Acknowledgement

In this draft, I would like to acknowledge Dr. Xiaolian Gao and colleagues at Department of Biology and Biochemistry, University of Houston, and Dr. Xiaochuan Zhou at LC Sciences, LLC for their assistance and inputs in peptide chip synthesis and analysis.

