## Supplementary figures and images for "A protein microarray-based *in vitro* transglutaminase assay platform for epitope mapping and immunogen design: implication for transglutaminase-mediated immunodominant determination"

### Synuclein TG array figure

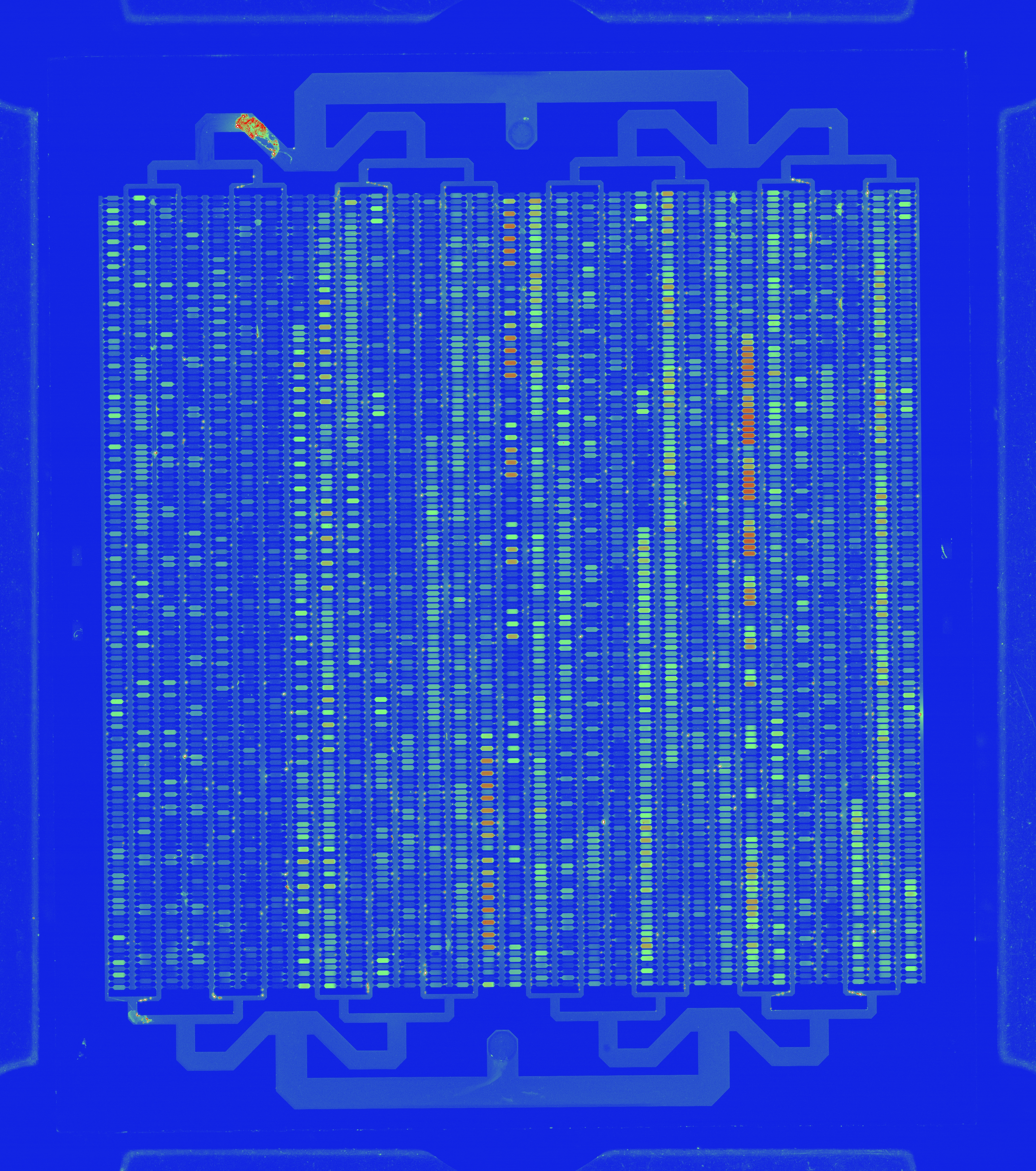
